# Comprehensive Evaluation of Protein Language Model Embeddings for Drug–Target Affinity Prediction

**DOI:** 10.64898/2026.08.31.748056

**Authors:** Matija Marijan, Ivan Tanasijević

## Abstract

Accurate identification of drug–target interactions is consequential for novel drug discovery and development. Deep learning methods for drug–target affinity (DTA) prediction have shown great promise in accelerating drug discovery and reducing development costs. Although graph neural networks have improved drug representation learning for DTA prediction tasks, many models still struggle to effectively and efficiently capture protein information, limiting overall prediction accuracy. In this work, we systematically evaluate the impact of pre-trained protein language models (PLMs) on the downstream task of predicting binding affinity between drugs and target proteins. We design multiple experiments across four different molecular representation backbones and assess the effect of incorporating PLM embeddings, comparing their performance to classical 1D convolution methods. We evaluate four families of PLMs which we integrate into PLM-GraphDTA, each built on distinct architectures and optimized for different tasks, including structure prediction, function prediction, and sequence unmasking. Additionally, we evaluate DeepGraphDTA, an architectural modification of the baseline convolution method designed to improve protein representation learning. The models are evaluated on two benchmark datasets, Davis and KIBA, using concordance index (CI) and mean squared error (MSE) as performance metrics. We further evaluate the generalization power of each model using cold-start train and test splits, and analyze the per-protein contribution to total CI. The results indicate simple architectural modifications to traditional convolution methods may be sufficient to bridge the gap to large pre-trained PLMs.

## 1 Introduction

Drug discovery is a complex and expensive process, in terms of both time and resources. In 2014, it was estimated that it costs approximately $2.8 billion over a span of 12.5 years to develop a single therapeutic [DiMasi et al., 2003, Paul et al., 2010, DiMasi et al., 2015]. A significant portion of the investment is allocated for the hit-to-lead and lead optimization phases of preclinical research [Hughes et al., 2011], where it is crucial to carefully curate libraries of candidate molecules. Despite advances and cost reductions in high-throughput screenings [Blay et al., 2020], these stages still benefit greatly from reliable computational estimates of drug–target affinity (DTA). Specifically, given that the vast majority of currently identified therapeutic targets are proteins [Bleicher et al., 2003], much of the research effort in this field focuses on estimating DTA between small molecules and target proteins.

An essential step in computational DTA evaluation is the reliable representation of both proteins and drugs, with minimal loss of critical information. Balancing representational power, computational costs, and data availability to achieve the desired predictive capability, in-silico DTA prediction methods span a spectrum of data- and physics-driven approaches [Zhang et al., 2024]. At one end of the spectrum, molecular docking methods estimate the binding free energy of drug–protein complexes by identifying the optimal three-dimensional (3D) conformation that minimizes the energy [Morris and Lim-Wilby, 2008]. Although this approach has been proven to provide reliable results [Ewing et al., 2001, Morris et al., 2009, Xu et al., 2021], it heavily relies on the high representational power of 3D crystal structures, which are often unavailable, and on free energy approximations that may accumulate errors at the level of detail necessary for reliable DTA estimation [Buttenschoen et al., 2024].

As an alternative, data-driven methods, led by early machine learning (ML) approaches, focus on simpler representations and overcome the aforementioned limitations by recognizing statistical patterns in the abundantly available data [Zhang et al., 2024]. One of the first robust and carefully validated models based on this paradigm was reported by Pahikkala et al. [2015], who introduced a regularized kernel model suited for DTA prediction. The kernel represents a separable similarity measure between two drug–target pairs as a product of the 2D molecular fingerprint similarity of the drugs and the sequence-based similarity measure of the proteins. Another important milestone for data-driven models was the development of SimBoost [He et al., 2017], a model based on gradient-boosted trees for DTA regression. Interestingly, its feature engineering relies heavily on handcrafted features of the drug–drug and target–target similarity networks, as well as the drug–target interaction network. Such features are easily captured by graph neural networks (GNNs), and SimBoost helped pave the way toward GNN-based DTA prediction.

With the advance of hardware-accelerated computing and the expansion of publicly available DTA databases, classical ML models and handcrafted features gave way to deep learning (DL) approaches [Pei et al., 2023, Xia et al., 2023, Yang et al., 2023, Qiu et al., 2024, Wu et al., 2024, Zhou et al., 2024]. The reduction in computational costs has enabled models to learn more meaningful representations of both drugs and proteins, directly from raw data. Öztürk et al. [2018] introduced a seminal DL model, DeepDTA. It utilizes convolutional neural networks (CNNs) to learn the representations and perform regression on the concatenated embeddings. Although the CNN approach to protein representation was a significant step-up compared to handcrafted features and similarity measures, DeepDTA processes drugs as SMILES strings, which, when combined with convolution, can lead to a loss of structural information. To address this limitation, Nguyen et al. [2021] introduced GraphDTA, a model that considers drugs as 2D molecular graphs and applies GNNs to learn structure-aware embeddings, with multiple GNN variants being explored.

Driven by the improvement enabled by the more sophisticated drug representation, recent literature has increasingly shifted focus toward improving protein representations [Jiang et al., 2020, Rifaioglu et al., 2021, Daga et al., 2023, Duy Nguyen and Son Hy, 2024]. Generally, representing proteins is a far more demanding task compared to drugs, not only because of the larger molecule size, but also because of the complex 3D structure, which is crucial for assessing interactions. Recent advances in deep learning and large language models (LLMs) have extended beyond natural language processing (NLP) into specialized domains such as protein analysis [Baek et al., 2021, Gligorijević et al., 2021, Jumper et al., 2021, Meier et al., 2021, Villegas-Morcillo et al., 2021, Lin et al., 2023, Yang et al., 2024, Hayes et al., 2025]. Protein language models (PLMs) treat amino acid sequences analogously to natural language, leveraging self-supervised learning and transformer architectures to capture both local and global dependencies, and generate informative embeddings for a wide range of predictive tasks, including structure and function prediction.

One example of a DTA model with improved protein representation, and a natural continuation of the previously mentioned DTA models, is PGraphDTA [Bal et al., 2023]. This model retains the GNN-based drug representation introduced in GraphDTA [Nguyen et al., 2021] and uses precomputed protein embeddings formulated by specialized LLMs.

In this work, we aim to reconcile and exhaustively assess CNN- and LLM-based protein representations, and focus on improving upon GraphDTA’s protein representation method, which we refer to as the baseline. We examine two strategies for protein representation that aim to improve the extraction of contextual, structural, and functional properties of proteins, addressing key limitations of prior methods. These strategies include modifying the convolutional architecture for learning protein embeddings to resemble the classical motif search approach [Sigrist et al., 2010], and leveraging representations from pretrained NLP models, specialized for protein structure and function analysis. We benchmark these approaches against state-of-the-art DTA prediction models and provide a comprehensive analysis of how different protein representations influence model performance and interpretability.

## 2 Methods

We describe the architecture of the DTA prediction framework in this section, which consists of several independent and interchangeable representation modules, suited for extensive evaluation and comparison experiments. The framework consists of two representation learning channels, as depicted in Figure 1: one for drugs and the other for proteins. Each channel uses specialized deep learning (DL) layers to extract information, producing two distinct latent representations of the input drug and protein. These representations are concatenated into a single vector and passed to a multi-layer perceptron (MLP) with two fully connected layers and a final regression head for binding affinity prediction.

**Figure 1:**
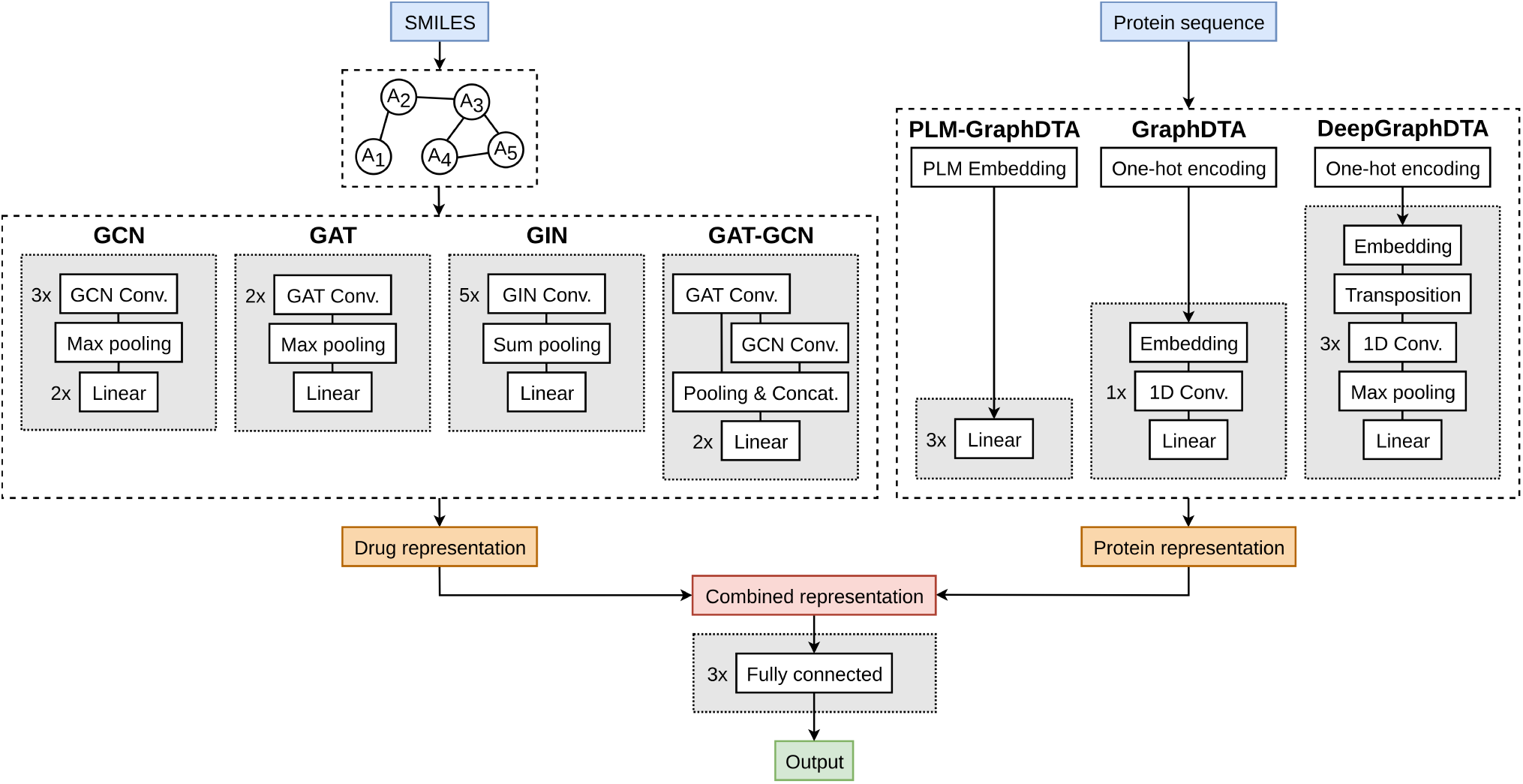
DTA protein representation evaluation framework. The top left section represents independent graph-based drug representation learning methods from GraphDTA (**GCN:** Graph convolution network, **GAT:** Graph attention network, **GIN:** Graph isomorphism network, **GAT-GCN:** Combined GAT-GCN network). On the right, three independent protein representation methods are shown (**PLM-GraphDTA:** Protein language model representations, **GraphDTA:** Baseline convolution, **DeepGraphDTA:** Transposed input convolution). Both channel outputs are combined and passed through the prediction head.

### 2.1 Drug representation

The drug-representation learning framework is based on GraphDTA [Nguyen et al., 2021], which portrays drugs as graphs of atomic interactions and applies GNN-based methods to learn graph-level representations. Drug graphs are constructed from SMILES strings [Weininger, 1988], where each node is assigned a feature vector that includes specific atom descriptors extracted using RDKit [Landrum et al., 2006]. The feature vectors, inspired by Ramsundar et al. [2019], include descriptors such as the atom symbol, the atom degree, the number of neighboring hydrogen atoms, the number of implicit hydrogen atoms on the central atom, and a flag indicating whether the atom is part of an aromatic structure.

Nguyen et al. [2021] proposed four distinct architectures for drug graph representation learning: graph convolutional networks (GCNs) [Kipf and Welling, 2016], graph attention networks (GAT) [Velicković et al., 2018], graph isomorphism networks (GIN) [Xu et al., 2018], and a combined GAT-GCN network.

The GCN-based model uses the standard graph convolutional operators introduced by Kipf and Welling [2016]. It consists of three graph convolutional layers followed by global max pooling to obtain a graph-level drug representation, which is finally processed by two fully connected layers.

The GAT-based model is based on graph attention networks [Velicković et al., 2018], which use attention mechanisms to weigh the contributions of neighboring nodes. It consists of two GAT layers, followed by global max pooling to obtain a graph-level representation, which is passed through a single fully connected layer.

The GIN-based model utilizes graph isomorphism operators [Xu et al., 2018]. It consists of five GIN blocks for learning node-level representations. Each block contains two linear layers followed by a GIN convolution layer. The learned node features are aggregated using global sum pooling, and a final linear layer produces the graph-level drug representation.

The GAT-GCN model combines graph attention and graph convolutional layers. It consists of a single GAT layer [Velicković et al., 2018], followed by a single GCN layer [Kipf and Welling, 2016]. Global max pooling is applied to the GAT output, while global mean pooling is applied to the GCN output. The pooled representations are concatenated and passed through two fully connected layers.

### 2.2 Protein representation

Traditionally, most protein representation methods are based on the primary structure, as amino acid sequences are readily accessible and easy to use. In contrast, graph-based representations are computationally complex and expensive, and tertiary structure data is often unavailable [Nguyen et al., 2021]. Integer encoding is commonly used to numerically represent sequences by assigning each amino acid a unique integer. Coupled with embedding layers, this approach enables DL models to learn meaningful protein representations.

We evaluate three protein representation approaches. Two methods follow the traditional encoding and embedding paradigm followed by different convolutional architectures, one of which is the baseline GraphDTA protein representation method. The other convolutional approach is DeepGraphDTA, which represents an architectural improvement to GraphDTA’s protein representation method. The third approach, which we refer to as PLM-GraphDTA, leverages precomputed PLM embeddings directly within the DTA prediction pipeline. Within it, we examine PLMs such as Evolutionary Scale Modeling (ESM) [Lin et al., 2023, ESM Team, 2024], Functional Residue Identification (DeepFRI) [Gligorijević et al., 2021], and Protein structure-sequence T5 (ProstT5) [Heinzinger et al., 2024].

All protein representation methods produce 128-dimensional protein representations, which are then concatenated with the drug representation to form a single 256-dimensional representation. For integer-encoded sequences, like Nguyen et al. [2021], we use a 128-dimensional embedding layer. The following sections provide a detailed overview of the three protein representation methods.

#### 2.2.1 GraphDTA—Baseline convolution

The baseline protein representation approach used in GraphDTA [Nguyen et al., 2021] uses a single 1D convolution layer to process protein embeddings. In this implementation, the convolution applies a filter to each amino acid embedding independently and aggregates the filtered information across all amino acids, treating each embedding as a separate input channel, as illustrated in Figure 2. A linear layer then transforms the output into the final latent representation of the input protein.

**Figure 2:**
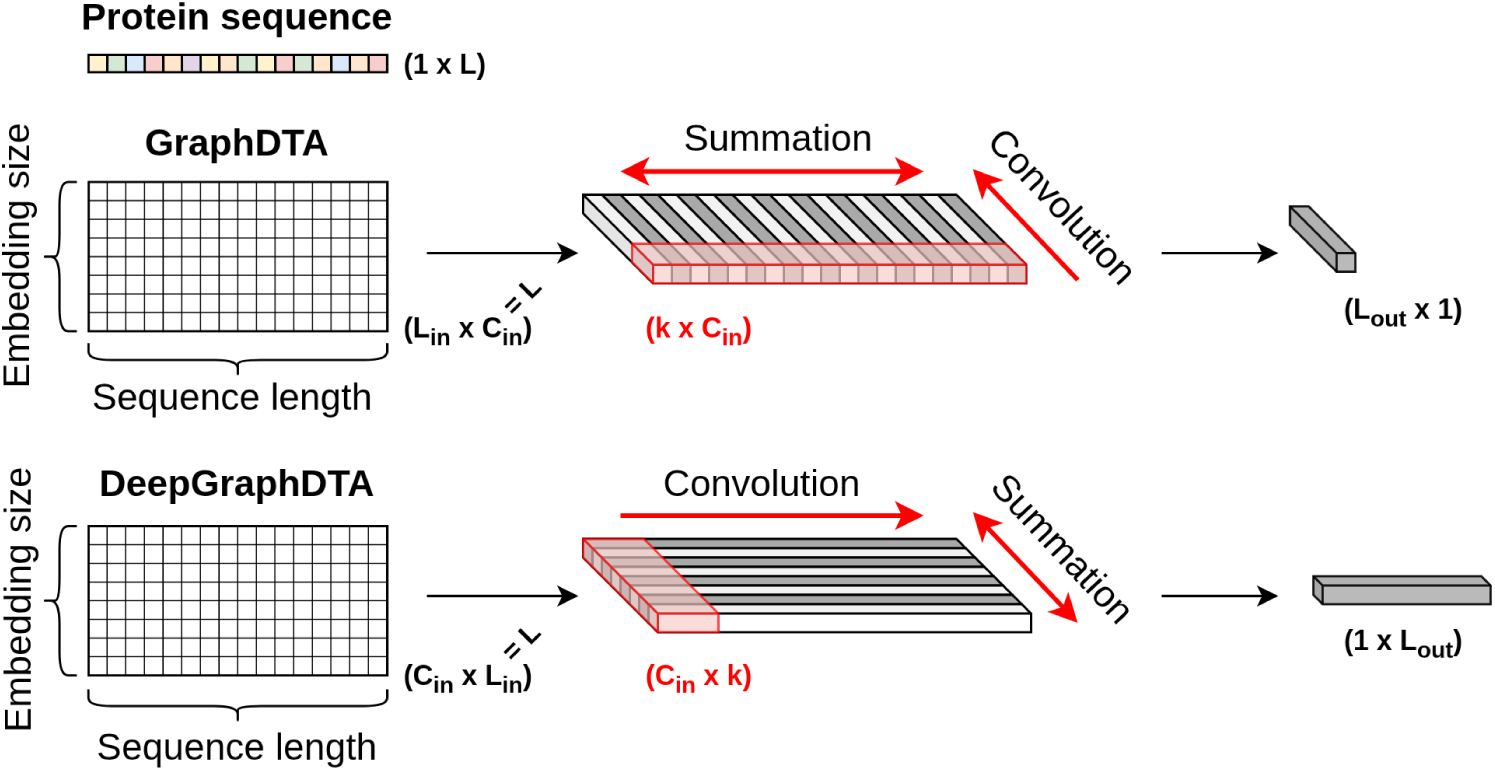
Different mechanisms of protein sequence 1D convolution, shown for a single filter. The baseline GraphDTA convolution is shown on top, and the DeepGraphDTA transposed input convolution is shown on the bottom. **L** represents the protein sequence length, **L_in_** and **L_out_** denote the input and output dimensions, respectively, **C_in_** represents the number of input channels, and **k** denotes the kernel size.

#### 2.2.2 DeepGraphDTA—Transposed input convolution

The DeepGraphDTA model transposes the input protein embedding before applying 1D convolution, enabling the filters to operate along the length of the sequence and aggregate the filtered information across all embedding dimensions, as depicted in Figure 2. This design ensures that sequential (temporal) dependencies are preserved, which may otherwise be lost. This modification aligns with the standard practice used in sequence-based CNNs, such as DeepDTA [Öztürk et al., 2018], and resembles the classical motif search approach [Sigrist et al., 2010], aiming to capture sequence patterns between neighboring amino acids. The model consists of three consecutive 1D convolution layers, followed by global max pooling and a linear layer for extracting the final protein representation.

#### 2.2.3 PLM-GraphDTA—Protein language model representations

To further investigate the impact of protein representations on DTA prediction, we leverage protein embeddings derived from pretrained PLMs designed for learning structural and functional properties of proteins, including ESM [Lin et al., 2023, ESM Team, 2024], DeepFRI [Gligorijević et al., 2021], and ProstT5 [Heinzinger et al., 2024]. ESM [Lin et al., 2023] is a transformer-based family of PLMs trained with masked language modeling on protein sequences without supervision, and specialized to predict the identity of randomly masked amino acids. This approach, combined with self-attention layers, enables the model to learn amino acid dependencies and capture both local and global structural and functional features. Trained on UniRef90 clusters [Suzek et al., 2015], ESM-2 allows the prediction of structural and functional properties and the computation of high-dimensional protein representations from input sequences. ESM Cambrian (ESMC) [ESM Team, 2024] is the successor to ESM-2, specifically trained to create expressive representations of protein sequences for downstream prediction, while lowering computational costs compared to previous models. ESMC is trained on sequences from UniRef90 [Suzek et al., 2015], MGnify [Richardson et al., 2023], and JGI [Nordberg et al., 2014].

The DeepFRI models [Gligorijević et al., 2021] are a family of protein function prediction models trained to predict Gene Ontology annotations [Ashburner et al., 2000] for each category: Molecular Function (MF), Biological Process (BP), and Cellular Component (CC); as well as Enzyme Commission (EC) numbers [Jeske et al., 2019]. The DeepFRI model explicitly incorporates structural information via *C_α_*–*C_α_* contact maps constructed from 3D structures retrieved from the Protein Data Bank [Berman et al., 2000], and uses a self-supervised language model with long short-term memory (LSTM) to extract features and learn protein representations. The LSTM model is pretrained on sequences from the Pfam database [Finn et al., 2014], and is utilized to extract residue-level features. A three-layer graph convolutional network then propagates structurally proximal features and constructs the final protein-level feature representation.

ProstT5 is an encoder-decoder, auto-regressive transformer model specialized for translating protein sequences into protein structures [Heinzinger et al., 2024], and trained with masked language modeling. It is a finetuned version of ProtT5-XL-U50 [Elnaggar et al., 2021], which was pretrained on UniRef50 [Suzek et al., 2015], and subsequently finetuned to translate between protein sequences and structures from AlphaFoldDB [Varadi et al., 2022]. Both ProstT5 and ProtT5-XL-U50 are based on the T5 transformer architecture [Raffel et al., 2020].

For PLM-based DTA prediction, we preprocess all protein sequences, generate protein representations, and cache them prior to training. During model training, the protein representations are passed directly to the DTA models and subsequently transformed through a linear layer to reduce dimensionality before performing affinity prediction.

## 3 Experiments

This section describes the experimental design for evaluating the proposed protein representation methods. We first introduce the benchmark datasets and evaluation metrics, followed by a description of the baseline models used for comparison. Next, we describe the experimental setup, including data splits, hyperparameter optimization, model selection, and holdout evaluation.

### 3.1 Datasets

We evaluate the DTA prediction models on two datasets widely used in previous work: the Davis kinase inhibitor selectivity dataset [Davis et al., 2011], and KIBA, a large-scale kinase inhibitory bioactivity dataset [Tang et al., 2014].

The datasets were obtained from the official GitHub repository for DeepDTA^1^, which provides the data in a format suitable for DTA prediction. The SMILES representations in the datasets were retrieved from PubChem [Kim et al., 2016], while the protein sequences were sourced from UniProt [Bateman et al., 2023] using their gene names^2^. Following Öztürk et al. [2018], all protein sequences are truncated or padded to a fixed length of 1000 residues.

#### 3.1.1 Davis dataset

The Davis dataset contains selectivity assays of the human kinase protein family, comprising 442 kinase proteins and 68 kinase inhibitors [Davis et al., 2011]. It reports the binding affinities, measured as the dissociation constant *K_d_*, for all 30 056 drug–target pairs. The *K_d_*values range from 0.016 nm to 10 000 nm, where higher values indicate weaker affinity. A *K_d_* of 10 000 nm is assigned to drug–target pairs with very weak or unobserved interactions. Notably, 20 931 pairs, more than half of the dataset, are reported with this maximum value.

Following the approach introduced by He et al. [2017], we applied a log transformation to the affinity values:

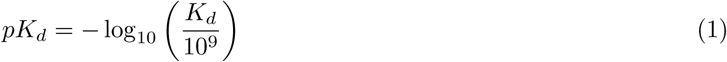

This transformation rescales the affinity range from 5.0, indicating weak affinity, to 10.8, the highest value.

#### 3.1.2 KIBA dataset

The KIBA dataset integrates kinase inhibitor bioactivity assays from multiple sources, including the dissociation constant *K_d_*, inhibitory constant *K_i_*, and the half-maximal inhibitory concentration *IC*_50_. These bioactivities are statistically adjusted and combined into KIBA scores, providing a unified measure of drug–target interaction strength [Tang et al., 2014].

The original KIBA dataset contains 52 498 ligands and 467 proteins, and a total of 246 088 KIBA scores. Following He et al. [2017], we use a filtered version that retains drug–target pairs with at least 10 interactions, resulting in 2111 drugs and 229 proteins. Unlike the Davis dataset, KIBA does not include all possible drug–target pairs; very weak or unmeasurable interactions are omitted, leading to a total of 118 254 reported interactions. The affinity values range from 0.0 to 17.2, with higher scores indicating stronger affinity.

### 3.2 Evaluation metrics

We evaluate the DTA prediction models using mean squared error (MSE) and concordance index (CI), following previous studies. MSE, a standard performance metric for regression tasks, is calculated as:

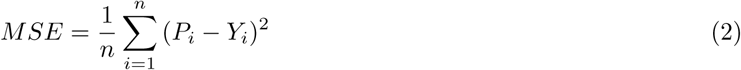

where *P_i_* are predictions, *Y_i_* are ground truths, and *n* is the number of samples. MSE was also used as the loss function for training the models.

While MSE is an adequate choice for both a differentiable loss function and an absolute performance metric, in affinity prediction scenarios, the quality of a predictive model is often better expressed by its discriminatory power and its ability to correctly rank affinities relative to one another. Therefore, we additionally evaluated performance using CI, a metric that quantifies the discriminatory power and predictive accuracy of nonlinear statistical models [Gönen and Heller, 2005], calculated as:

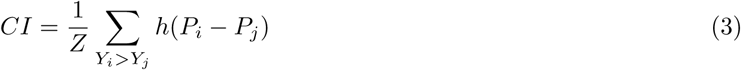

where *P_i_* and *P_j_* are predicted values for an input pair with true affinity values *Y_i_* and *Y_j_*, such that *Y_i_ > Y_j_*, *Z* is a normalization constant, and *h*(*x*) is a step function returning 1, 0.5, and 0 for *x >* 0, *x* = 0, and *x <* 0, respectively [Pahikkala et al., 2015]. This metric measures how well a model preserves the relative order of predicted values, estimating the probability that, for a randomly chosen input pair, the order of the predicted values is concordant with their true order [Harrell Jr et al., 1984].

### 3.3 Baseline models

We primarily base the comparison against GraphDTA [Nguyen et al., 2021], which is described in Sections 2.1 and 2.2.1. We also compare our proposed models with state-of-the-art models KronRLS [Pahikkala et al., 2015], SimBoost [He et al., 2017], DeepDTA [Öztürk et al., 2018], and PGraphDTA [Bal et al., 2023].

KronRLS [Pahikkala et al., 2015] is a regression algorithm that implements Kronecker regularized least squares to predict drug–target interactions. It utilizes Kronecker kernel functions to model pairwise similarities between drugs and targets, integrating various similarity measures. Drug features are calculated based on encoded 3D structural similarity coefficients, while protein feature vectors are derived from Smith-Waterman (S-W) sequence alignment scores [Smith et al., 1981]. The Kronecker product of these similarity matrices forms a comprehensive kernel that captures the interactions between all drug–target pairs, enabling accurate affinity prediction.

SimBoost [He et al., 2017] predicts DTA by leveraging similarity-based features and gradient boosted regression trees. It builds on the KronRLS method [Pahikkala et al., 2015] by incorporating three types of features: the properties of individual drugs and targets based on similarity values, network-based features derived from drug and target similarity graphs, and interaction-based features from affinity-weighted drug–target networks.

DeepDTA [Öztürk et al., 2018] employs convolutional neural networks (CNNs) to learn latent representations from protein sequences and drug SMILES. It uses integer encoding to enable 1D convolution on drug SMILES representations, and adopts a protein feature extraction approach similar to the DeepGraphDTA model described in Section 2.2.2. The learned representations are concatenated and passed through fully connected layers for affinity prediction.

PGraphDTA [Bal et al., 2023] is a model that, similarly to PLM-GraphDTA, utilizes PLM embeddings for protein representations, and evaluates multiple PLMs, including ProtTrans [Elnaggar et al., 2021], DistilProtBERT [Geffen et al., 2022], SeqVec [Heinzinger et al., 2019], and ESM-2 [Lin et al., 2023]. In addition, Bal et al. [2023] explore the integration of protein contact maps through their PGraphDTA-CM variants. To ensure a controlled and fair comparison with PLM-GraphDTA, we restrict our analysis to the ESM-2-based variant that does not incorporate contact maps. In contrast to PLM-GraphDTA, PGraphDTA uses only the GAT-based drug encoder from GraphDTA [Nguyen et al., 2021], and the graph-level drug representations are constructed by applying both max pooling and learnable weighted sum pooling to node-level representations and concatenating the resulting vectors. Furthermore, PGraphDTA uses higher-dimensional latent representations of both drugs and proteins relative to PLM-GraphDTA.

### 3.4 Experiment setup

This section details the experiment setup, including data splits, hyperparameter optimization, model selection, and strategies for evaluating the proposed protein representation methods within the DTA prediction pipeline.

To ensure a fair comparison during cross-validation and final evaluation, we used the same predefined training and testing sets of drug–target pairs as proposed by Öztürk et al. [2018], and adopted in subsequent studies. The train–test split follows a 5:1 ratio.

All experiments were conducted under identical software and hardware conditions, on a server node allocated with an AMD EPYC 7742 CPU with 64 cores, 128 GB RAM, and an NVIDIA A100 GPU (40 GB VRAM). Details of the software environment used for developing, training, and evaluating the models are available at the official GitHub repository for PLM-GraphDTA^3^.

The training hyperparameters are summarized in Table 1, and they are consistent with those used in GraphDTA [Nguyen et al., 2021]. The same hyperparameters are used for all other experiments, including 5-fold cross-validation and cold-start experiments.

**Table 1:**
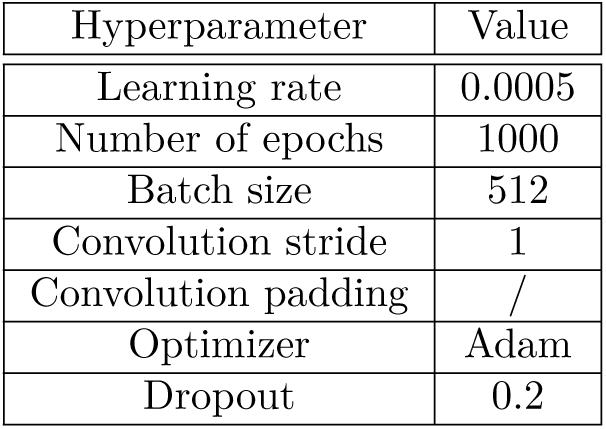
Hyperparameters used for DTA prediction model training.

In all proposed models, ReLU activations follow each convolutional and intermediate dense layer, with batch normalization applied to stabilize training. Dropout is applied after dense layers to reduce overfitting. The number of trainable and frozen parameters in all model variants is listed in the appendix B.

#### 3.4.1 DeepGraphDTA hyperparameter optimization

We performed 5-fold cross-validation (CV) to optimize the 1D convolution kernel size for protein representations in DeepGraphDTA, using a grid search over the range [8, 16, 24, 32]. For each of the 4 fixed molecular graph representation methods and 2 datasets, the optimal kernel size was selected based on the highest mean CI on the validation folds, with lower standard deviation (SD) and MSE as tie-breakers. Detailed results of the DeepGraphDTA hyperparameter optimization experiment are shown in appendix C.

#### 3.4.2 PLM-GraphDTA model selection

Similarly, we conducted 5-fold CV to determine the optimal PLM for protein representations in PLM-GraphDTA. We compared and evaluated several models: three ESM2 model sizes (8M, 150M, and 650M), two ESMC model sizes (300M and 600M), four ontology-specific DeepFRI models (BP, CC, EC, and MF), and ProstT5. For each of the 4 molecular representations and 2 datasets, the best-performing model was selected based on the highest mean CI, with SD and MSE as tie-breakers. Detailed results of the PLM-GraphDTA model selection experiment are shown in appendix D.

The pretrained ESM-2 and DeepFRI models are publicly available at their respective official GitHub repositories^4^, while the pretrained ProstT5 and ESMC models are available at their respective official HuggingFace repositories^5^.

#### 3.4.3 Final evaluation

Using the determined best model architectures, we performed final model training on all 5 train folds, and evaluated their performance on the holdout test fold. Final model evaluation was performed and averaged for 5 different random seeds for reproducibility. We report the average MSE and CI scores, and compare the results from DeepGraphDTA and PLM-GraphDTA against all baseline models. Furthermore, we use paired t-tests to determine if the improvements in CI using DeepGraphDTA and PLM-GraphDTA are statistically significant in relation to GraphDTA, and finally, we report the results of the paired t-tests to determine if there exists a statistically significant improvement in CI using PLM-GraphDTA over DeepGraphDTA.

#### 3.4.4 Cold-start evaluation

A known limitation of DTA prediction models is the lack of diverse DTA data, which often leads to poor generalizability. We conduct three different experiments, each specialized for evaluating the generalizability of prediction models. Specifically, we perform drug cold-start training and evaluation, in which there exists a holdout set of drugs unseen in the training set. Similarly, we follow up with protein cold-start experiments, where a set of protein sequences is held out from the training set. Finally, we perform fully blind cold-start evaluation, where a set of drug–target pairs is held out entirely from the training set, making this scenario the most difficult.

Each of the three experiment scenarios is prepared by performing a 5:1 train-test data split, where either a random set of drugs/proteins is held out (along with all of their accompanying input pairs), or a random set of drug–protein pairs is held out. We perform five random data splits, with 5 different random initial seeds. Afterwards, we train each of the model variants on the train set, and report the MSE and CI on the test set. We report the mean MSE and mean CI across the five different data splits.

## 4 Results and discussion

Figure 3 compares the MSE and CI metrics of the original GraphDTA protein representation, and the optimal DeepGraphDTA and PLM-GraphDTA variants chosen through 5-fold CV for each of the 4 fixed molecular graph representations on the holdout test fold of the Davis and KIBA datasets. DeepGraphDTA and PLM-GraphDTA models show statistically significant improvements in CI compared to the baseline GraphDTA for all molecular representations, on both datasets, with all p-values *<* 0.05. Although PLM-GraphDTA outperforms DeepGraphDTA in almost every scenario by both mean CI and MSE, results of the paired t-test indicate a statistically significant improvement in CI in using PLM-GraphDTA over DeepGraphDTA only for the GAT-GCN protein representation, paired with the ProstT5 PLM, with a p-value of 0.049. These results indicate that using precomputed language may not lead to significant improvements in performance compared to a relatively small architectural improvement of the baseline GraphDTA protein representation. A conclusion might be made that the use of PLMs may unnecessarily increase computational complexity and costs, with no major improvements to performance.

**Figure 3:**
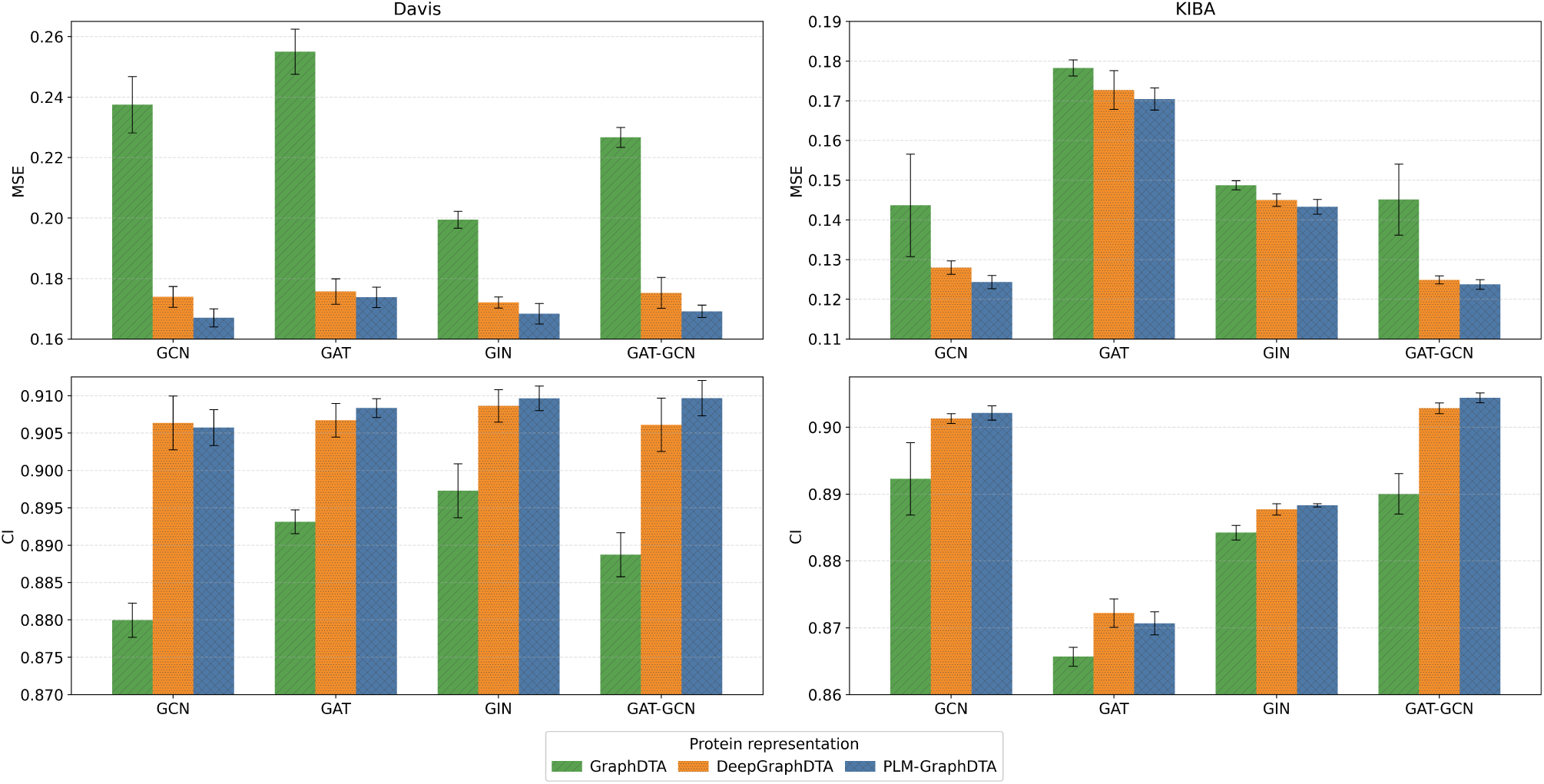
Comparison of protein representation methods on the Davis (left) and KIBA datasets (right).

Tables 2 and 3 compare the MSE and CI performance metrics of DeepGraphDTA and PLM-GraphDTA models, with five baselines on the holdout test fold on the Davis and KIBA datasets, respectively.

**Table 2:** Affinity prediction performance comparison on the Davis dataset, sorted by ascending CI.

| Method | Protein representation | Drug representation | CI ↑ | MSE ↓ |
| --- | --- | --- | --- | --- |
| Baseline models |  |  |  |  |
| KronRLS [Pahikkala et al., 2015] | Similarity vector | Similarity vector | 0.870 | 0.359 |
| GraphDTA [Nguyen et al., 2021] | 1D conv. | GCN | 0.880 | 0.237 |
| SimBoost [He et al., 2017] | Similarity features | Similarity features | 0.885 | 0.246 |
| GraphDTA [Nguyen et al., 2021] | 1D conv. | GAT-GCN | 0.889 | 0.227 |
| GraphDTA [Nguyen et al., 2021] | 1D conv. | GAT | 0.893 | 0.255 |
| GraphDTA [Nguyen et al., 2021] | 1D conv. | GIN | 0.897 | *0.199* |
| PGraphDTA [Bal et al., 2023] | ESM2 650M | GAT | 0.898 | 0.206 |
| DeepDTA [Öztürk et al., 2018] | 1D conv. | 1D conv. | *0.902* | 0.224 |
| Proposed models |  |  |  |  |
| PLM-GraphDTA | ESMC 600M | GCN | **0.906** | **0.167** |
| DeepGraphDTA | 1D conv. (8) | GAT-GCN | **0.906** | **0.175** |
| DeepGraphDTA | 1D conv. (16) | GCN | **0.906** | **0.174** |
| DeepGraphDTA | 1D conv. (16) | GAT | **0.907** | **0.176** |
| PLM-GraphDTA | ESMC 600M | GAT | **0.908** | **0.174** |
| DeepGraphDTA | 1D conv. (32) | GIN | **0.909** | **0.172** |
| PLM-GraphDTA | ESMC 600M | GIN | **0.910** | **0.168** |
| PLM-GraphDTA | ESMC 600M | GAT-GCN | **0.910** | **0.169** |

*Note:* Values in italics represent the best for the baseline models, while values in bold represent values better than baseline. The values in brackets represent the convolution kernel size selected by 5-fold CV.

**Table 3:** Affinity prediction performance comparison on the KIBA dataset, sorted by ascending CI.

| Method | Protein representation | Drug representation | CI ↑ | MSE ↓ |
| --- | --- | --- | --- | --- |
| Baseline models |  |  |  |  |
| KronRLS [Pahikkala et al., 2015] | Similarity vector | Similarity vector | 0.782 | 0.411 |
| SimBoost [He et al., 2017] | Similarity features | Similarity features | 0.836 | 0.222 |
| DeepDTA [Öztürk et al., 2018] | 1D conv. | 1D conv. | 0.863 | 0.194 |
| GraphDTA [Nguyen et al., 2021] | 1D conv. | GAT | 0.866 | 0.178 |
| PGraphDTA [Bal et al., 2023] | ESM2 650M | GAT | 0.872 | 0.160 |
| GraphDTA [Nguyen et al., 2021] | 1D conv. | GIN | 0.884 | 0.149 |
| GraphDTA [Nguyen et al., 2021] | 1D conv. | GAT-GCN | 0.890 | 0.145 |
| GraphDTA [Nguyen et al., 2021] | 1D conv. | GCN | *0.892* | *0.144* |
| Proposed models |  |  |  |  |
| PLM-GraphDTA | ProstT5 | GAT | 0.871 | 0.170 |
| DeepGraphDTA | 1D conv. (24) | GAT | 0.872 | 0.173 |
| DeepGraphDTA | 1D conv. (16) | GIN | 0.888 | 0.145 |
| PLM-GraphDTA | ESMC 600M | GIN | 0.888 | **0.143** |
| DeepGraphDTA | 1D conv. (8) | GCN | **0.901** | **0.128** |
| PLM-GraphDTA | ProstT5 | GCN | **0.902** | **0.124** |
| DeepGraphDTA | 1D conv. (32) | GAT-GCN | **0.903** | **0.125** |
| PLM-GraphDTA | ProstT5 | GAT-GCN | **0.904** | **0.124** |

*Note:* Values in italics represent the best for the baseline models, while values in bold represent values better than baseline. The values in brackets represent the convolution kernel size selected by 5-fold CV.

The results show clear performance improvements in both CI and MSE with DeepGraphDTA and PLM-GraphDTA in comparison with the baseline models on the Davis dataset. These improvements may indicate improved abstraction power for capturing essential protein properties and describing drug–target interactions in comparison to GraphDTA. In comparison to other baseline methods, the results confirm the results reported by Nguyen et al. [2021], which is that GNNs significantly improve the representational power of DTA prediction models.

The PLM-GraphDTA model with an ESMC 600M PLM paired with the GCN drug representation relatively improves MSE by 16.08% in comparison to the best baseline MSE, achieved by GraphDTA for the GIN drug representation. The improvements in CI are less pronounced, with PLM-GraphDTA with an ESMC 600M paired with the GAT-GCN drug representation achieves a relative improvement of 0.89% in comparison with the best baseline CI, achieved by DeepDTA.

On the KIBA dataset, the results show clear performance improvements in both CI and MSE with both DeepGraphDTA and PLM-GraphDTA for the GCN and GAT-GCN molecular representations, in comparison with the baseline models. On the other hand, the GAT and GIN representations do not provide significant improvements to the best baseline. The PLM-GraphDTA model with the ProstT5 and GAT-GCN molecular representation achieves a 1.35% improvement in CI and a 13.89% improvement in MSE relative to the best baseline, which is GraphDTA paired with a GCN for drug representations.

The performance gains are more pronounced with MSE, rather than CI, which may suggest that current methods are reaching a performance plateau in DTA prediction, and deeper and more expressive model architectures may not improve prediction accuracy greatly. On the other hand, an equally likely explanation might be that most of the current DTA datasets are too small and not too diverse, which may lead to overfitting to those datasets. This is also evident when analyzing the generalizability of DTA prediction models.

### 4.1 Cold-start evaluation results

Figure 4 and 5 show the results for the cold-start evaluation experiments. Immediatelly noticable is the degradation in both MSE and CI performance metrics, across all data splits and datasets, along with much more unstable values, with large SDs. The models roughly preserve their relative ranking across all scenarios, with few exceptions. Notably, the GIN GraphDTA variant outperformed the GIN DeepGraphDTA variant in the drug cold-start scenario on the Davis dataset, and the GAT GraphDTA variant outperformed the GAT DeepGraphDTA variant in the drug cold-start scenario on the KIBA dataset.

**Figure 4:**
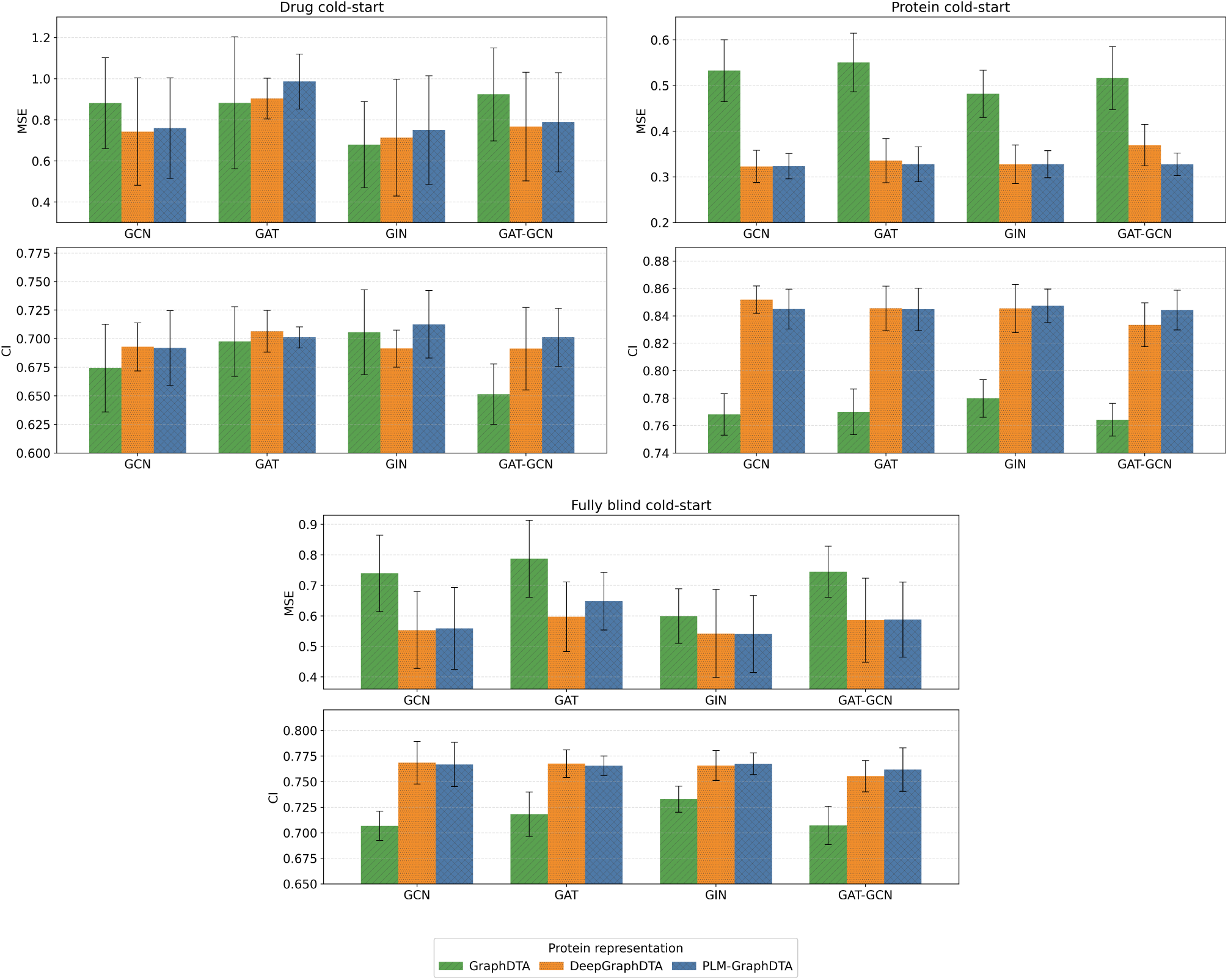
Cold-start evaluation on the Davis dataset (drug cold-start – top left, protein cold-start – top right, fully blind cold-start – bottom).

**Figure 5:**
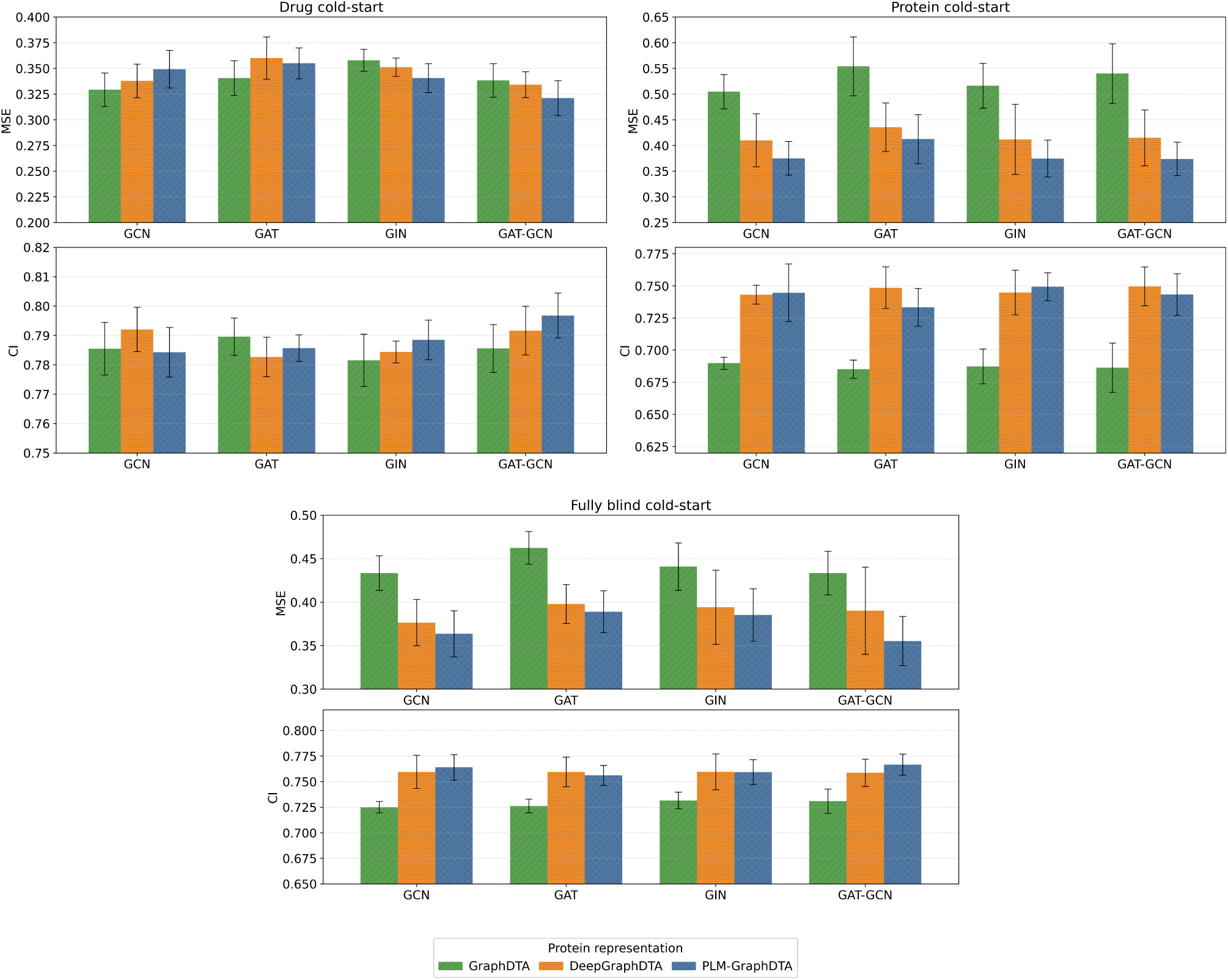
Cold-start evaluation on the KIBA dataset (drug cold-start – top left, protein cold-start – top right, fully blind cold-start – bottom).

**Figure 6:**
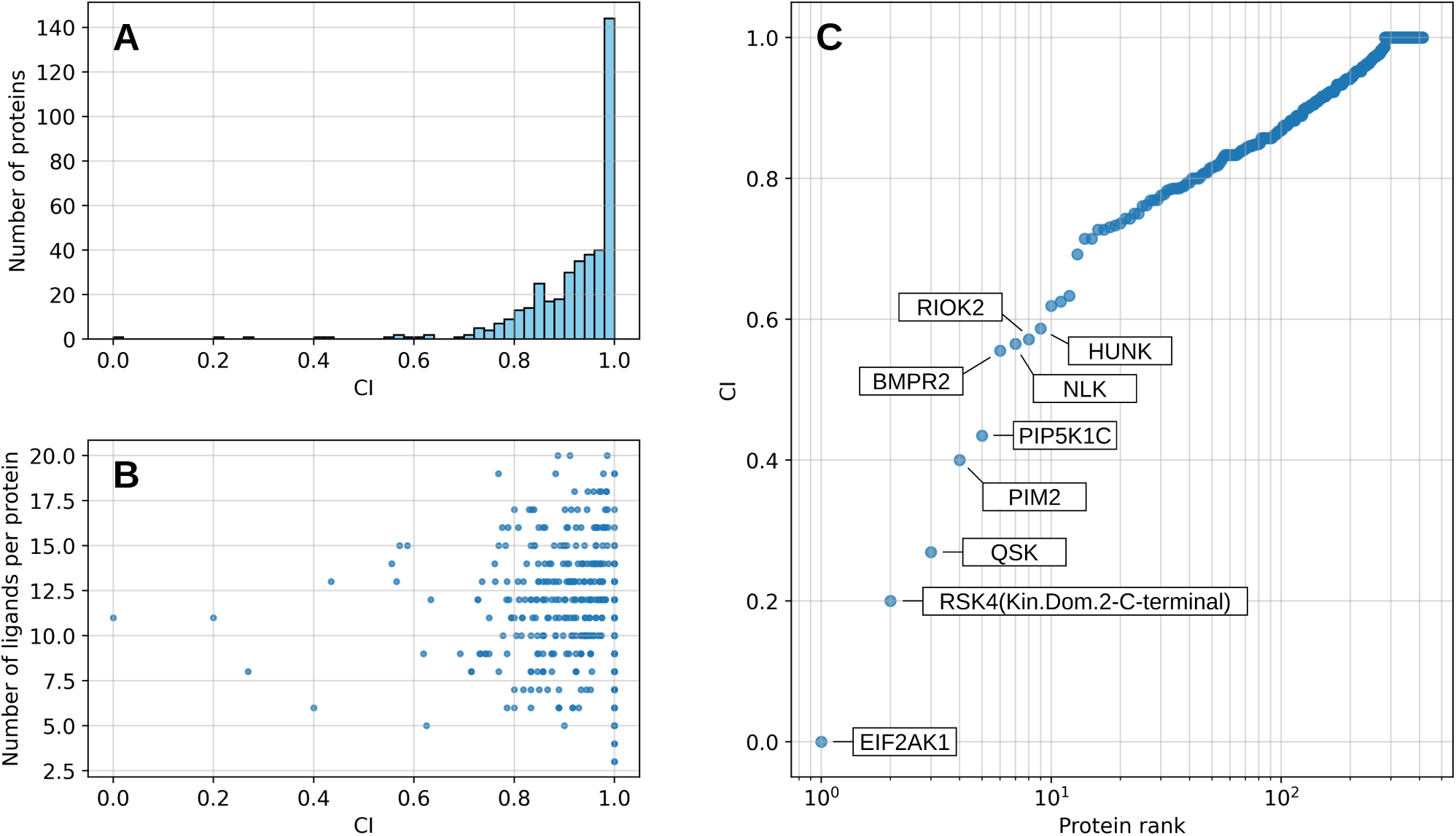
PLM-GraphDTA (ESMC 600M, GIN) CI analysis on the Davis dataset. **A:** Histogram of CI values in relation to number of proteins. **B:** Distribution of CI values in relation to number of ligands per specific protein. **C:** Protein ranking by CI values. The top 10 worst-performing proteins are indicated by their HGNC gene names.

Notably, the improvements introduced with enhanced protein representations with DeepGraphDTA and PLM-GraphDTA become almost indiscernible in the drug cold-start scenarios on both datasets. This might lead to a conclusion that molecular representations still carry the grunt of the burden in DTA representation problems. On the other hand, in fully blind cold-start scenarios, DeepGraphDTA and PLM-GraphDTA are significantly more performant than GraphDTA, even though a certain set of drugs is fully held out from the training set, similarly to the drug cold-start scenario.

The GAT-GCN molecular representation is the most stable in both datasets in all scenarios, likely due to its high parameter count. This might indicate that further research into more complex, and deeper molecular representation methods is warranted.

## 5 Analysis

Following the prediction error analysis by Nguyen et al. [2021], in this section, we perform a similar analysis and take a deeper look into the predicted affinity values that the proposed models output. In this section, we focus on identifying if any proteins contribute disproportionately to mean CI. We analyze the histogram of the CI values in relation to number of proteins in the dataset, the distribution of CI values in relation to the number of ligands per specific protein, and finally, we take a look at the worst-performing proteins in relation to the CI value for that protein.

We conduct these analyses for the models that have shown best and most stable performance in 5 fold CV. We focus on PLM-GraphDTA with the ESMC 600M PLM and the GIN drug representation for the Davis dataset, and PLM-GraphDTA with the ProstT5 PLM and the GAT-GCN network for the KIBA dataset.

Through these results, the differences between the two datasets come to light more clearly. The Davis dataset contains almost 6 times more protein sequences than drugs, and has a large number of data pairs which have the lowest output affinity. This likely leads to a larger number of affinities being correctly ranked per protein, irregardless of the number of drugs present in the testing set. On the contrary, the KIBA dataset contains much more drugs than protein sequences, leading to CI values being much more similar to a normal distribution, albeit with a peak at *CI* = 1. At this value, unsurprisingly, we see that no proteins have more than 25 paired drugs in the testing set. In the KIBA dataset, all of the worst-performing proteins have a relatively small number of attributed drugs in the testing set, whereas in the Davis dataset, although the worst-performing proteins have a lower number of attributed drugs in the testing set than in the KIBA dataset, the number is proportional to the average number of ligands per protein in the Davis dataset. This result might contribute to the conclusion that specific proteins (and their accompanying drugs) contain difficult interaction patterns which are difficult to represent within the latent space, leading to higher errors in DTA prediction scenarios.

## 6 Conclusion

In this paper, we exhaustively investigate the impact of protein representation methods on drug–target affinity prediction and propose modifications to these methods to improve prediction accuracy. Building on the drug representations introduced with GraphDTA [Nguyen et al., 2021], we focus our analysis on the impact of protein representations, focusing on redesigned convolutional layers with DeepGraphDTA, and integrating precomputed protein representations derived from pretrained protein language models through PLM-GraphDTA.

We benchmarked these approaches with state-of-the-art DTA prediction models using the Davis [Davis et al., 2011] and KIBA [Tang et al., 2014] datasets, and show that the testend protein representations improve performance with statistically significant improvements in both concordance index and mean squared error. The improvements still hold in more difficult cold-start scenarios, indicating improved abstraction and representation power in comparison to the baselines.

The key findings include that a simple architectural modification to the baseline 1D convolution method achieves performance comparable to large pretrained PLMs, suggesting that CNN architectures may be sufficient for many DTA prediction tasks, eliminating the need for the computational overhead of language models. Furthermore, while the PLM-based representations generally outperform both baseline and DeepGraphDTA models, the improvements are often not statistically significant.

The choice of molecular representation remains crucial, as seen in cold-start experiments, where a handful of held out drugs may sway the overall performance of the models greatly. Future work should not exclude drug representation research and development, especially in cold-start scenarios. Also, the severe degradation in cold-start scenarios indicate and highlight the continued need for more diverse training data. We also identified and corrected significant data quality issues in the commonly-used Davis dataset, where mutated proteins were incorrectly represented, affecting over 14% of protein sequences, and subsequently, a number of publications on DTA prediction.

Despite these performance improvements and findings, our analyses reveal that DTA prediction may be reaching a performance plateau with current architectures, and more specifically, current datasets. The modest improvements in CI across models and datasets, combined with poor generalization in cold-start scenarios, suggest fundamental limitations that cannot be overcome solely through enhanced protein representations.

Future research may benefit from integrating complementary protein representations and structural features, including pocket information, contact maps, and docking simulations [Yang et al., 2022, Yazdani-Jahromi et al., 2022, Wu et al., 2024]. Attention mechanisms and LSTMs have shown promise in capturing key drug–target dependencies while maintaining efficiency [Lin et al., 2022, Zhao et al., 2022, Pei et al., 2023]. A deeper comparison of models using PLM embeddings [Daga et al., 2023, Duy Nguyen and Son Hy, 2024] could further guide optimization. Finally, recent advances in 3D protein structure prediction [Baek et al., 2021, Jumper et al., 2021, Lin et al., 2023] open up new opportunities for graph-based representations, though reducing model complexity remains a key challenge. On another note, future work could evaluate the potential performance gains of implementing an ensembling approach to both drug and protein representations. Additionally, expanding dataset diversity and size, as well as developing more robust evaluation protocols that better assess real-world generalization capabilities, are critical directions for advancing the field.

## Author contributions

**Matija Marijan:** Data curation, Formal analysis, Investigation, Methodology, Software, Validation, Writing – original draft, Writing – review & editing. **Ivan Tanasijević:**Conceptualization, Supervision, Writing – review & editing.

## Acknowledgments

The authors thank Jelena Těsić for her assistance with data acquisition and analysis, and Dejan Miřcetić for his valuable feedback on the manuscript.

## Availability of data and materials

All source codes, trained models, and datasets are publicly available at https://github.com/matija-marijan/PLM-GraphDTA.

^†^This work was conducted while the authors were affiliated with the Institute for Artificial Intelligence Research and Development of Serbia.

^1^https://github.com/hkmztrk/DeepDTA/tree/master/dat^a^

^2^An issue we identified concerning protein mutations in the Davis dataset is examined and explained in appendix A.

^3^https://github.com/matija-marijan/PLM-GraphDT^A^

^4^https://github.com/facebookresearch/es^m^; https://github.com/flatironinstitute/DeepFRI

^5^https://huggingface.co/EvolutionaryScal^e^; https://huggingface.co/EvolutionaryScale

## **A** Sequence mutations in the Davis dataset

We discovered that the preprocessed version of the Davis dataset commonly used for DTA prediction models contains only 379 unique protein sequences, rather than the 442 originally reported. The remaining 63 nonunique sequences correspond to mutated proteins examined in the original study by Davis et al. [2011], in which drug–target affinities for various mutants were measured to investigate their impact on binding. These mutations are indicated in the dataset by distinct gene descriptors, which typically specify amino acid substitutions. For example, ABL1(E255K) indicates that the glutamic acid at position 255 in the parent protein ABL1 is replaced by lysine.

However, when preparing datasets for deep learning-based DTA prediction, an oversight can easily occur when retrieving sequences from UniProt [Bateman et al., 2023] using these gene descriptors. Since UniProt does not typically include such mutated proteins as separate entries, it returns the parent protein sequence instead. Consequently, entries that should represent distinct protein variants share the same sequence in the dataset. This affects more than 14% of the protein entries and results in identical sequences being associated with different binding affinities for the same drug, making it difficult for models to learn meaningful drug–target relationships. As a result, this inconsistency propagates through many DTA prediction models that evaluate their performance on this version of the Davis dataset.

A paper on DTA prediction by Monteiro et al. [2022] provides a corrected version of the Davis dataset, in which most mutations have been resolved, although without detailed documentation. In the corrected dataset, the only remaining non-unique sequences correspond to nonstandard mutation types, such as phosphorylated and non-phosphorylated proteins, the FLT3 protein and its internal tandem duplication variant FLT3(ITD), and the CDK4-cyclin D1 and CDK4-cyclin D3 protein variants.

Phosphorylation regulates protein functions by adding phosphate groups at sites distinct from the active site [Ubersax and Ferrell Jr, 2007]. Although it can affect protein conformation, it does not alter the protein’s primary structure, making it difficult to distinguish between variants. Internal tandem duplication of the FLT3 gene, reported in acute myeloid leukaemia patients [Gilliland and Griffin, 2002], does not follow a predictable pattern, which can lead to variations in the resulting protein sequence. Furthermore, to the best of our knowledge, the CDK4-cyclin D1 and CDK4-cyclin D3 complexes are not available in UniProt. Only the individual CDK4, cyclin D1, and cyclin D3 proteins are available. We retained these entries as they appear in the original dataset.

After additional processing, which includes filling missing sequences, we trained all models and report prediction results on this corrected version of the Davis dataset. Accounting for mutated/duplicated protein sequences reduces MSE by approximately 15% in comparison to the original version of the Davis dataset.

## **B** Parameter count

The number of trainable parameters for each protein representation model is listed in Table 4, along with the number of frozen parameters in the pretrained language models. The number of trainable parameters in each convolution-based protein representation method is listed in Table 5. The number of trainable parameters in each molecular GraphDTA representation is listed in Table 6. The number of trainable parameters in the prediction head is identical in all molecule–protein representation method combinations, and it is equal to 525 825 trainable parameters.

**Table 4:**
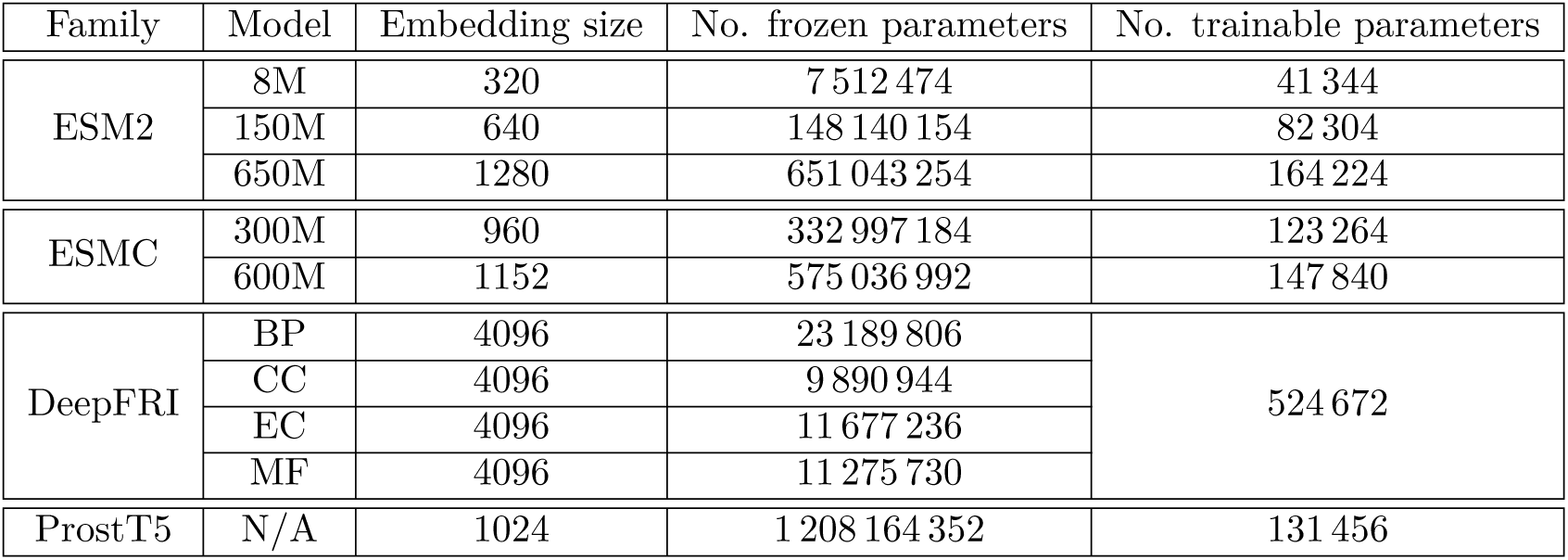
Number of parameters in PLM-GraphDTA protein representation models.

| Family | Model | Embedding size | No. frozen parameters | No. trainable parameters |
| --- | --- | --- | --- | --- |
| ESM2 | 8M | 320 | 7 512 474 | 41 344 |
|  | 150M | 640 | 148 140 154 | 82 304 |
|  | 650M | 1280 | 651 043 254 | 164 224 |
| ESMC | 300M | 960 | 332 997 184 | 123 264 |
|  | 600M | 1152 | 575 036 992 | 147 840 |
| DeepFRI | BP | 4096 | 23 189 806 | 524 672 |
|  | CC | 4096 | 9 890 944 |  |
|  | EC | 4096 | 11 677 236 |  |
|  | MF | 4096 | 11 275 730 |  |
| ProstT5 | N/A | 1024 | 1 208 164 352 | 131 456 |

**Table 5:** Number of parameters in convolution-based protein representation models.

| Model | 1D conv. layers | Kernel size | No. trainable parameters |
| --- | --- | --- | --- |
| GraphDTA | [32] | 8 | 755 104 |
| DeepGraphDTA | [32, 64, 96] | 8 | 114 880 |
|  |  | 16 | 213 184 |
|  |  | 24 | 311 488 |
|  |  | 32 | 409 792 |

**Table 6:** Number of trainable parameters in GraphDTA molecular representation models.

| Model | No. trainable parameters |
| --- | --- |
| GCN | 515 182 |
| GAT | 179 916 |
| GIN | 16 576 |
| GAT-GCN | 3 205 988 |

## **C** DeepGraphDTA hyperparameter optimization results

In this section, we present the results of the DeepGraphDTA hyperparameter optimization 5-fold CV experiments. Figures 8 and 9 show the CV performance metrics on the Davis and KIBA datasets for the 4 different protein 1D convolution kernel sizes within DeepGraphDTA.

**Figure 7:**
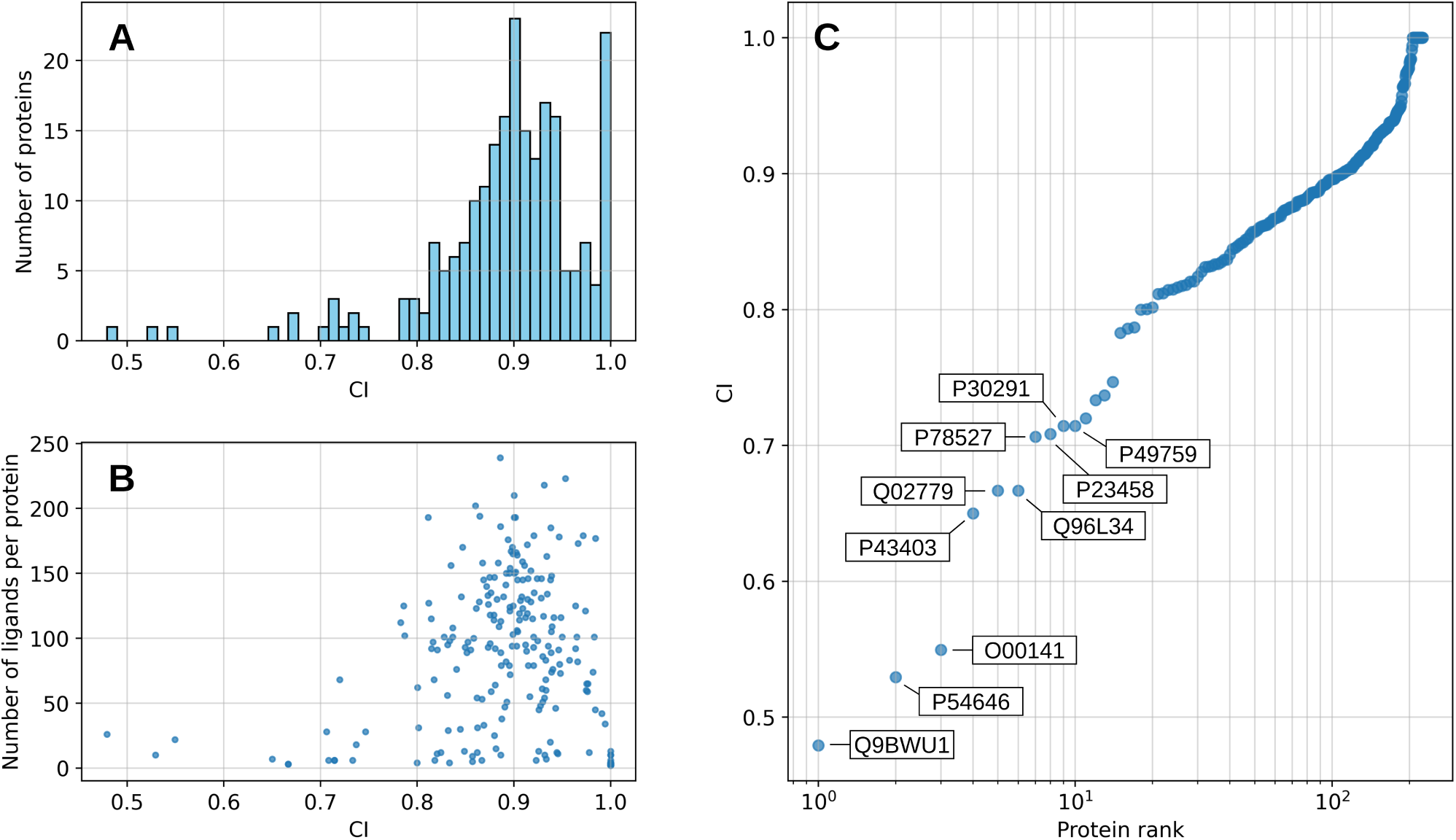
PLM-GraphDTA (ProstT5, GAT-GCN) CI analysis on the KIBA dataset. **A:** Histogram of CI values in relation to number of proteins. **B:** Distribution of CI values in relation to number of ligands per specific protein. **C:** Protein ranking by CI values. The top 10 worst-performing proteins are indicated by their HGNC gene names.

**Figure 8:**
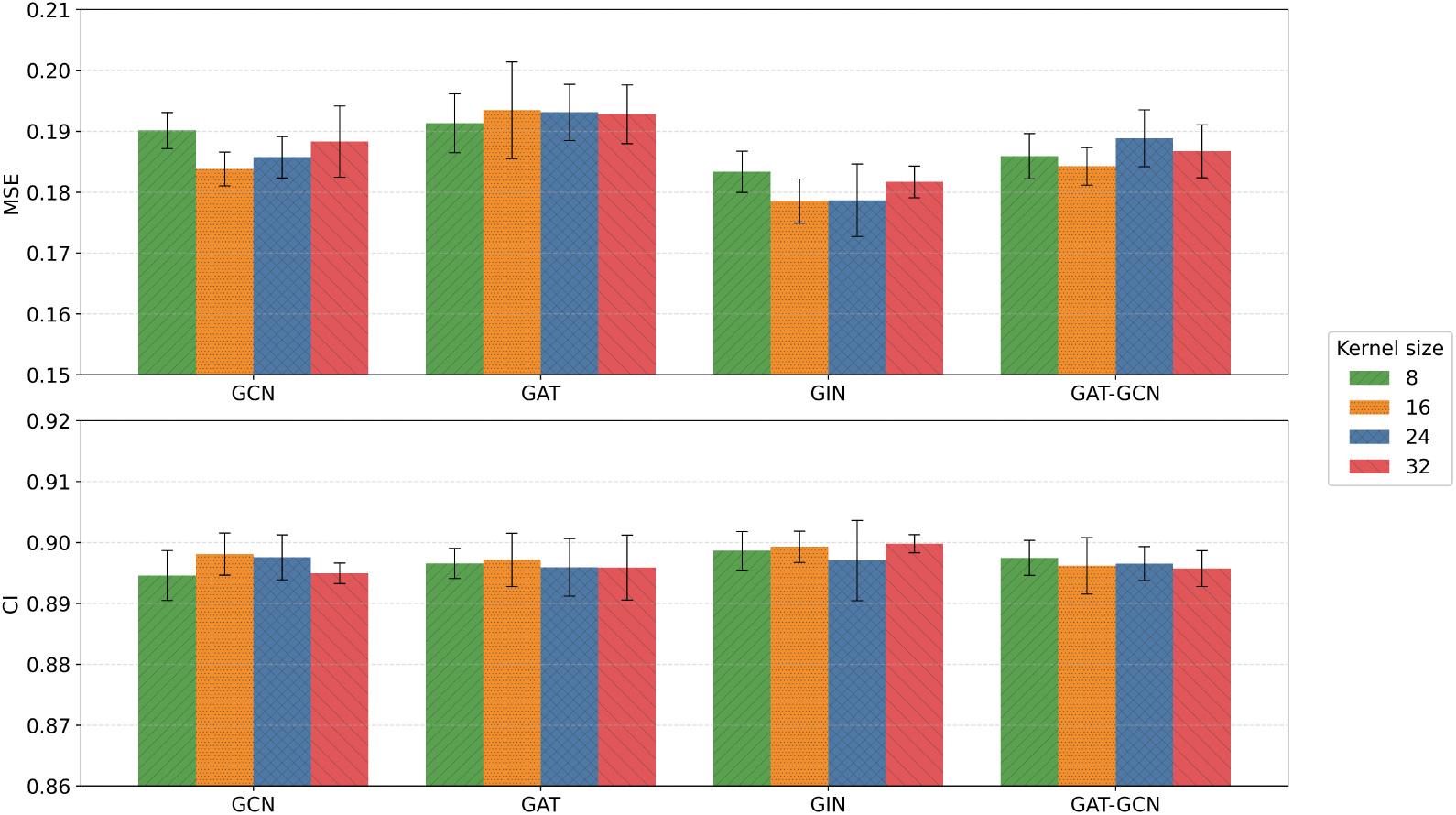
Protein 1D convolution kernel size 5-fold CV results on the Davis dataset.

**Figure 9:**
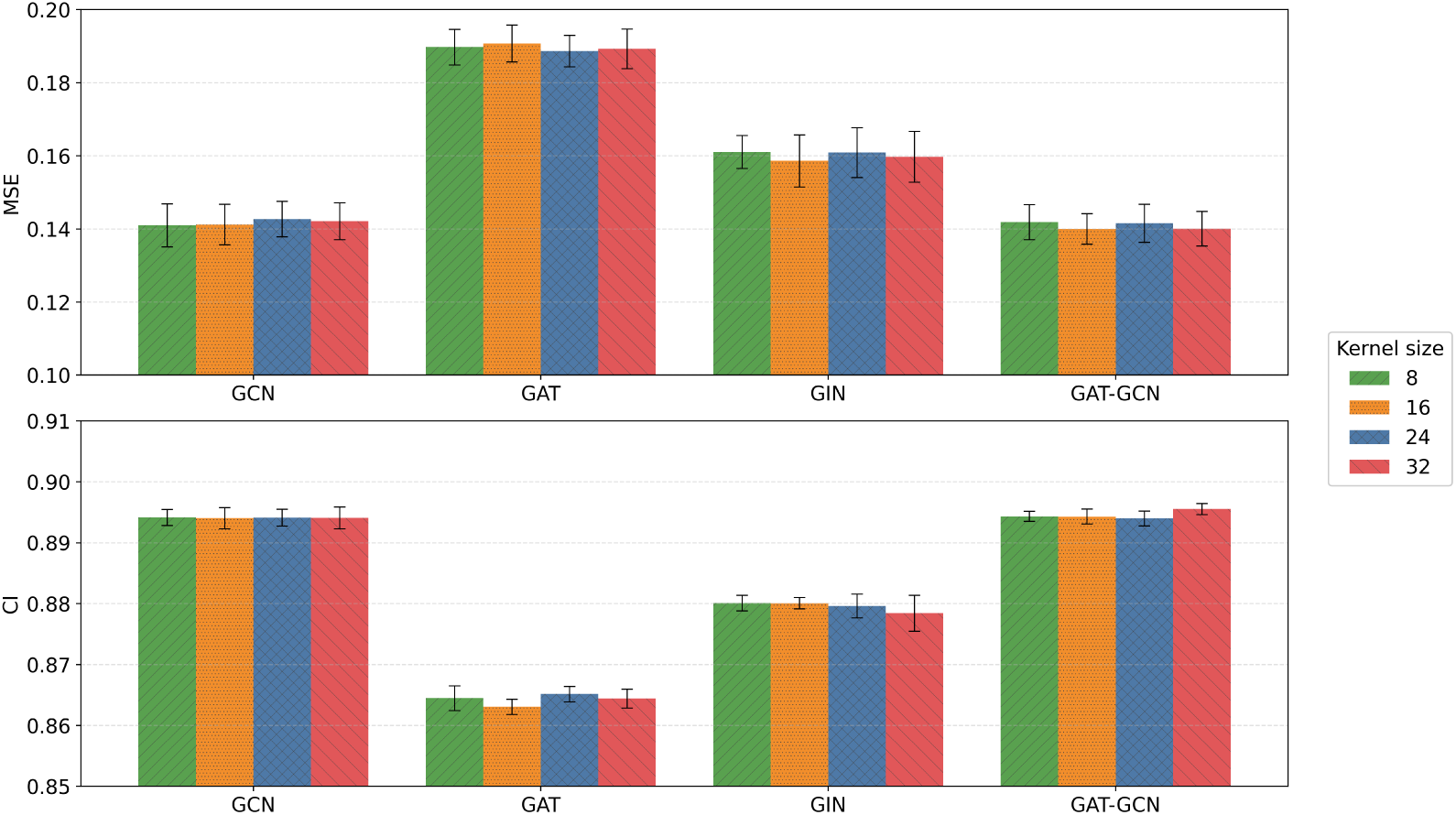
Protein 1D convolution kernel size 5-fold CV results on the KIBA dataset.

As evident in the figures, no specific pattern can be identified in the results. This might indicate that the specific kernel size might not be as important as the number of total convolution layers and channels. In this study, we only focused on the kernel size, whereas future work could focus on extensively investigating how CNN depth and number of convolution channels affect protein representations in downstream tasks.

## **D** PLM-GraphDTA model selection results

In this section, we present the results of the 5-fold CV experiments for determining the strongest PLM for the different molecular representations within PLM-GraphDTA. Figures 10 and 11 show the MSE and CI performance metrics for the ESM-2, ESMC, DeepFRI, and ProstT5 protein representations within PLM-GraphDTA, for all 4 molecular representations (GCN, GAT, GIN, and GAT-GCN).

**Figure 10:**
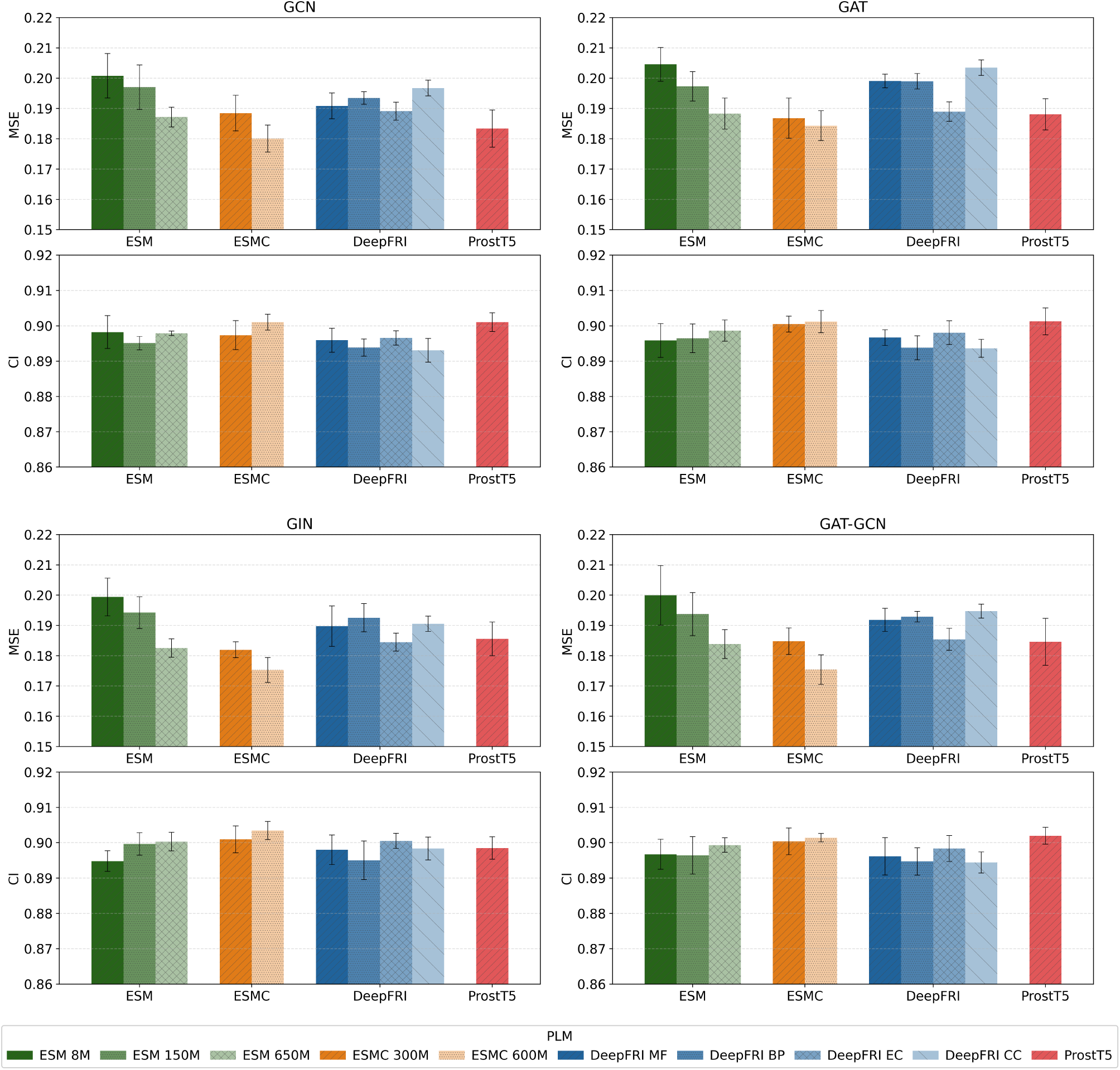
PLM 5-fold CV results for different molecular representations on the Davis dataset (GCN – top left, GAT – top right, GIN – bottom left, GAT-GCN – bottom right).

**Figure 11:**
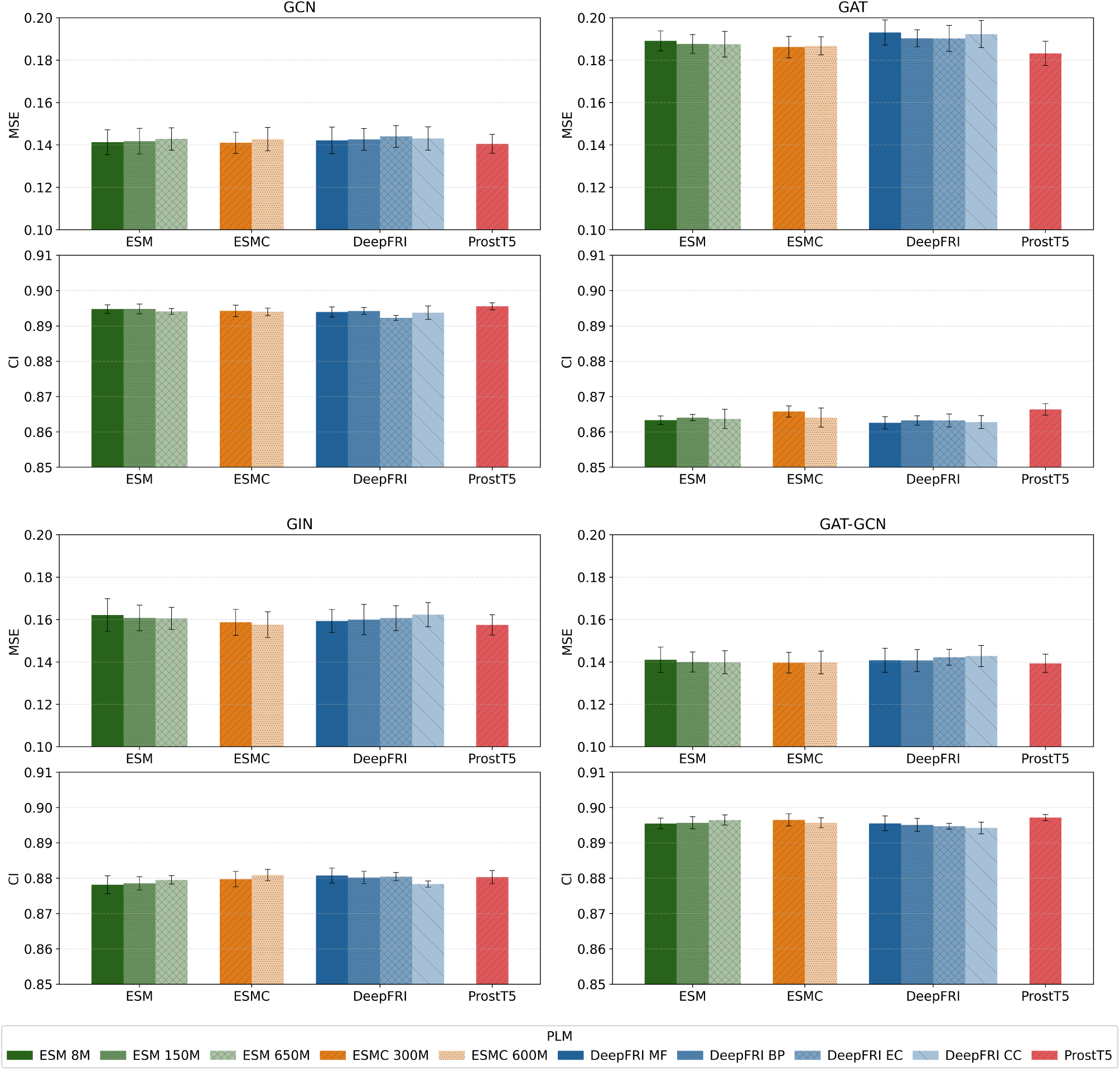
PLM 5-fold CV results for different molecular representations on the KIBA dataset (GCN – top left, GAT – top right, GIN – bottom left, GAT-GCN – bottom right).

Unsurprisingly, the larger the PLM, the better the performance of the downstream PLM-GraphDTA model, with few exceptions. The ESMC models consistently outperform the other PLMs on the Davis dataset, whereas the differences in MSE and CI are much more subtle on the KIBA datasets, with ProstT5 and the ESMC 600M proving strongest on the KIBA dataset.

